# RXFP1 expression is regulated by miR-144-3p in Fibroblasts from Patients with Idiopathic Pulmonary Fibrosis

**DOI:** 10.1101/159459

**Authors:** Harinath Bahudhanapati, Jiangning Tan, Justin A Dutta, Stephen B Strock, Yingze Zhang, Daniel J Kass

## Abstract

Relaxin has been considered as a potential therapy for patients with pulmonary fibrosis. We have previously shown, however, that a potential limitation of relaxin-based therapy for Idiopathic Pulmonary Fibrosis (IPF) is the loss of expression of the relaxin receptor Relaxin/Insulin Like Receptor 1 (RXFP1) expression in fibroblasts. The molecular mechanism for RXFP1 down-regulation in IPF patients remains unclear. To determine whether microRNAs play a role in RXFP1 gene expression, we employed a bioinformatics approach to identify microRNAs (miRs) that are predicted to target RXFP1. By *in silico* analysis, we identified a putative target site in the RXFP1 mRNA for the miR-144 family. We found that miR-144-3p was upregulated in IPF fibroblasts compared to donor lung fibroblast controls. Forced miR-144-3p mimic expression reduced RXFP1 mRNA and protein levels and increased expression of the myofibroblast marker alpha-smooth muscle actin (α-SMA) in donor lung fibroblasts. IPF lung fibroblasts transfected with a miR-144-3p inhibitor increased RXFP1 expression and reduced α-SMA expression. A lentiviral luciferase reporter vector carrying the WT 3’UTR of RXFP1 was repressed more in lung fibroblasts whereas vector carrying a mutated miR-144-3p binding site exhibited less sensitivity to endogenous miR-144-3p expression, suggesting that RXFP1 is a direct target of miR-144-3p. Thus, miR-144-3p is highly expressed in IPF fibroblasts and acts as a negative regulator of RXFP1 protein expression.

---

Idiopathic Pulmonary Fibrosis (IPF) is a chronic, debilitating, and progressive scarring of the lung parenchyma which ultimately compromises gas exchange and progresses to respiratory failure and death in many by four years after the diagnosis (1,2). The hallmark of fibrosis, which is shared by all organs, is the dysregulated, and seemingly unending, architectural destruction caused by the accumulation of activated fibroblasts and the deposition of extracellular matrix. Multiple lines of
evidence have suggested that the hormone relaxin is a potentially powerful inhibitor of fibrosis (3,4). These include the seminal observation that mice genetically engineered to lose expression of the hormone relaxin (encoded in humans by RLN2) developed a progressive, age-related multi-organ fibrosis that was reversible with exogenous relaxin 4–6). In further support of the importance of this pathway, we have shown that gene expression levels for the relaxin receptor, RXFP1, in IPF is directly associated with pulmonary function (7). Furthermore, RXFP1 protein expression was dramatically decreased in both IPF lungs and IPF lung fibroblasts compared to donor controls. Because the loss of RXFP1 expression in IPF may desensitize fibroblasts from the positive effects of relaxin-like agonists (7), a strategy to increase RXFP1 expression in IPF might prove to be an effective therapeutic approach.

Little is known, however, about the transcriptional regulation of RXFP1 expression. Certain endocrine signals have been associated with RXFP1 expression. For example, estrogen and progesterone treatment increase RXFP1 expression in ligamentous tissue from ovariectomized female rats (8). Alpha, but not β2 adrenergic stimulation of cardiac myocytes increases expression of RXFP1 (9). We and others have shown TGFβ stimulation can lead to decreased RXFP1 expression (7,10). Further highlighting the heterogeneity of fibrosis in different organs, increased RXFP1 expression has been observed in liver fibrosis (11,12). To our knowledge, there are no data on the regulation of RXFP1 by particular transcription factors.

One relatively unexplored mechanism to regulate RXFP1 expression is the role of microRNAs. MicroRNAs are noncoding small RNAs, about 22 nucleotides in length, that can bind to the 3’ UTR of target genes to repress their translation and/or induce degradation of target gene mRNA by incomplete base pairing. Dysregulation of miRNAs such as miR-29 (13), miR-21 (14), miR-155 (15), let-7d (16), miR-30 (17), and miR-133a (18) has been implicated in the pathogenesis of pulmonary fibrosis. Only synthetic microRNAs have been associated with regulation of RXFP1 expression (19). In this study, we tested the hypothesis that dysregulation of microRNA expression in IPF fibroblasts regulates RXFP1 gene expression.

## RESULTS

*miR-144-3p is upregulated in lung fibroblasts from IPF patients*—We have previously shown that loss of RXFP1 at the level of mRNA in IPF is associated with more impaired pulmonary function in patients. To understand potential mechanisms that may regulate RXFP1 transcription in lung fibroblasts, the principal effector cells in fibrosis, we employed several public database prediction programs to identify potential microRNA species that may regulate RXFP1 gene expression. Using the miRanda (microrna.org) and Targetscan 7.0 online databases, we identified a putative miR-144-3p targeting site in the 3’UTR of human RXFP1 mRNA (Fig 1*A*). While miR-144-3p is highly conserved in primates and lower vertebrates, the RXFP1 target site for miR-144-3p is conserved only among human, chimpanzee, rhesus, bovine and rat genomes. MiR-144-3p is predicted to have strong homology with the 3’UTR of RXFP1 mRNA. We chose miR-144-3p for further testing.

**FIGURE 1.**
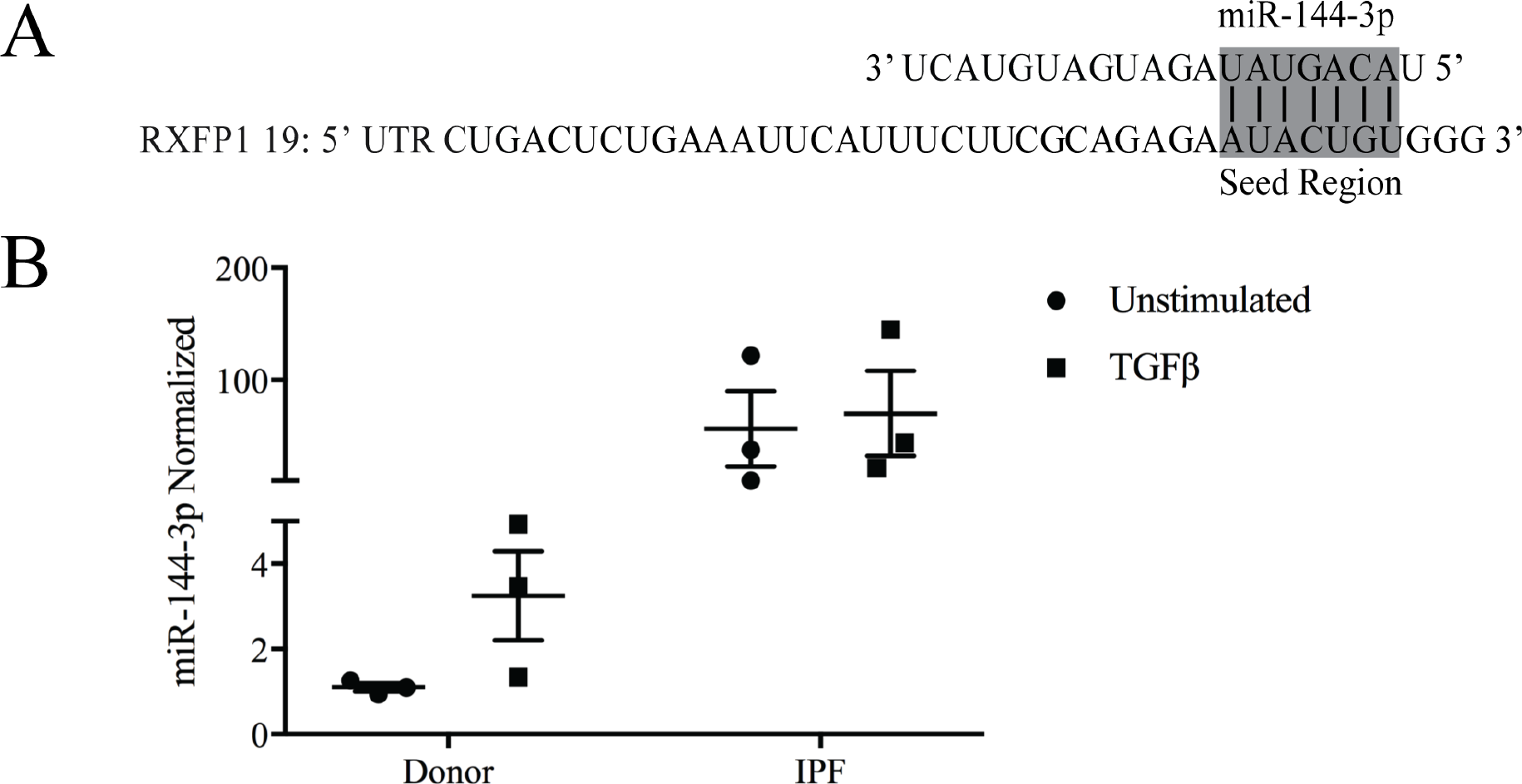
miR-144-3p expression is elevated in IPF fibroblasts. (*A*) The seed region of miR-144-3p that targets the 3’ UTR of RXFP1.(*B*) Donor and IPF lung fibroblasts were stimulated with and without TGFβ, and RNA was isolated for qPCR. Significantly more miR- 144-3p was detected in IPF lung fibroblasts compared to donor controls. Significant effects of disease were detected (p<0.05) by two-way ANOVA.

First we determined the levels of miR-144-3p in IPF and donor lung fibroblasts using quantitative RT-PCR with and without TGFβ stimulation. We found that fibroblasts derived from IPF lungs showed greater than 50-fold higher miR-144-3p compared to donor lung fibroblast controls (p<0.05 for significant effects of disease origin, by two-way ANOVA, N=3). TGFβ stimulation increased expression of miR-144-3p in donor lung fibroblasts but not in IPF fibroblasts (Fig 1*B*).

*Human RXFP1 mRNA is a direct target of miR-144-3p*—Next, we determined whether miR-144-3p overexpression in Donor and IPF lung fibroblasts can downregulate RXFP1 mRNA and protein levels. We transfected donor and IPF lung fibroblasts with 10 nM miR-144-3p mimic and processed these cells for quantitative RT-PCR and protein detection of RXFP1 (Fig 2). We found by qRT-PCR that donor lung fibroblasts have higher basal levels of RXFP1 mRNA compared to IPF lung fibroblasts (p<0.0016, by two-way ANOVA followed by post-hoc testing), confirming our previous findings (7) (Fig 2*A*). Second, qRT-PCR results showed that RXFP1 is significantly downregulated at mRNA level in donor lung fibroblasts following miR-144-3p overexpression (p<0.0001, by two-way ANOVA followed by post-hoc testing) (Fig 2*A*). These data support the notion that miR-144-3p regulates RXFP1 expression. For specificity of the effect of miR-144-3p on expression of RXFP1, we determined the effect on other relaxin receptor genes RXFP2 and RXFP3. *In silico*, miR-144-3p isnot predicted to target RXFP2 or RXFP3. Expression of human RXFP1 orthologue RXFP2 was unaffected by miR-144-3p overexpression compared to control mimic based on qRT-PCR data. Expression of RXFP3 was undetectable in donor or IPF lung fibroblasts (Fig 2*B*). RXFP3 is known to be expressed in central nervous system, but it is not known to be expressed in lungs (20).

**FIGURE 2.**
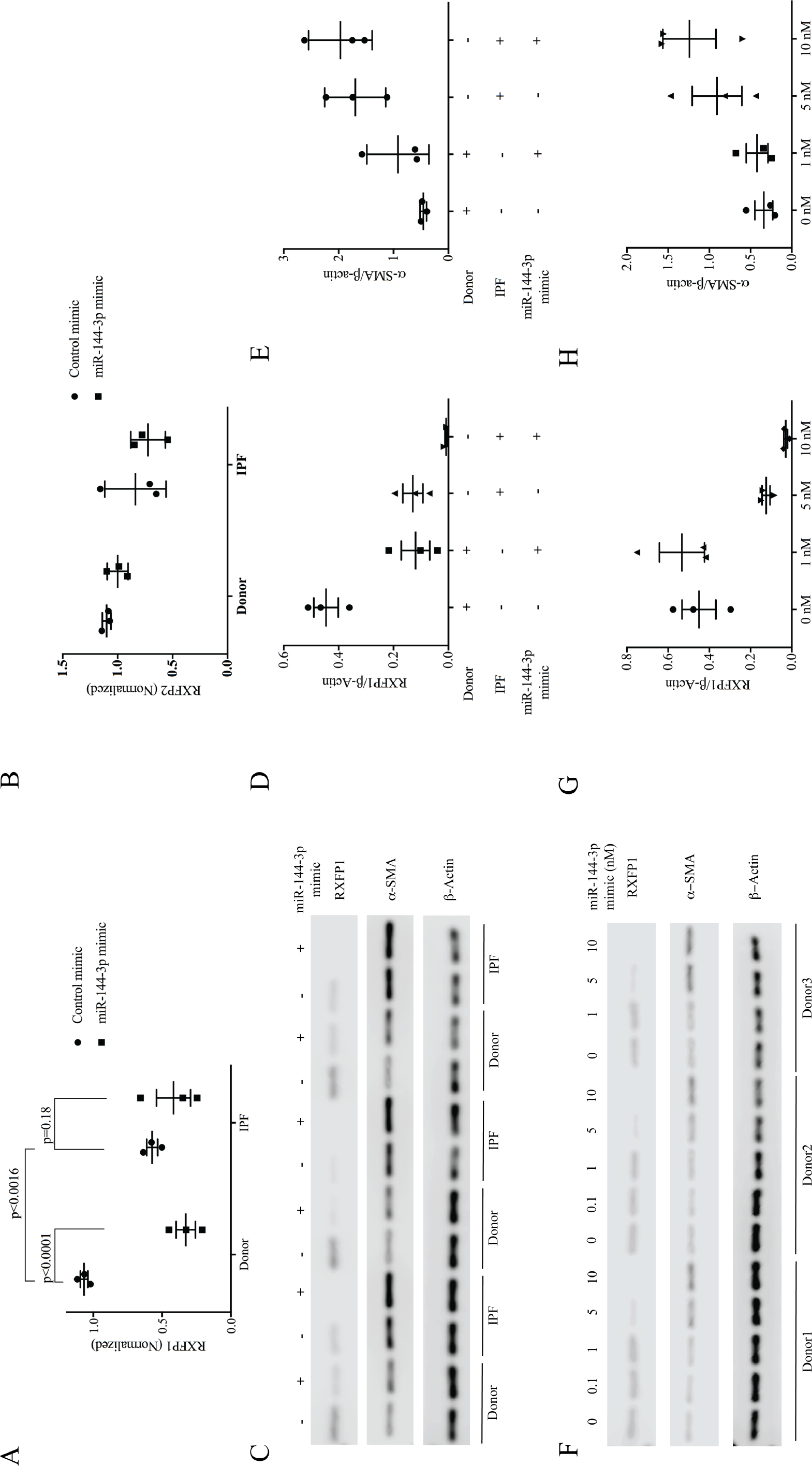
RXFP1 mRNA is a direct target of miR-144-3p. *(A)* Donor and IPF lung fibroblasts were treated with miR-144-3p mimic or scrambled control. RNA was isolated and processed for qPCR for RXFP1. Significantly higher basal levels of RXFP1 mRNA were detected in donor fibroblasts. Treatment of donor fibroblasts with the miR-144-3p significantly decreased RXFP1 expression (p<0.0016, by two-way ANOVA. N=3). Data are expressed as mean ± S.E.M, normalized to the internal standard control gene PPIA (B) miR-144-3p mimic had no effect on expression levels of RXFP2 both in fibroblasts from lungs of IPF patients or from donor lungs. RXFP3 levels were undetectable in both donor and IPF lung fibroblasts (*C*) Donor and IPF lung fibroblasts were transfected with 10 nM miR-144-3p mimic or control. miR-144-3p mimic significantly decreased RXFP1 expression in both donor and IPF lung fibroblasts whereas a-SMA expression significantly increased in IPF lung fibroblasts. Samples were analyzed by Western Blotting. Signals were analyzed with the C-DIGIT imager, quantified with Image Studio^™^ Software and normalized to β-actin (*D*) Image Studio^™^ quantification of band intensity of RXFP1 were normalized to β-actin. Data are expressed as mean ± S.E.M., normalized to β-actin. Significant effects of tissue origin (IPF *v* Donor, p= 0.0006, by two-way ANOVA, N=3) and miR-144-3p transfection (IPF *v* Donor, p= 0.0006, by two-way ANOVA, N=3) on RXFP1 expression were observed. Significant interaction between tissue origin and miR-144-3p transfection was detected (p<0.03). (*E*) Image Studio^™^ quantification of band intensity of α-SMA were normalized to β-actin. Significant effects of tissue origin (IPF *v* Donor, p=0.0038, by two-way ANOVA, N=3) on aSMA expression were detected. No significant interaction was detected. (*F* Donor lung fibroblasts were transfected with increasing concentration of miR-144-3p (0.1, 1, 5, and 10 nM). miR-144-3p mimic decreased the expression of RXFP1 and upregulated the expression of α-SMA in a dose-dependent manner. (*G*) Image Studio^™^ quantification of band intensities of RXFP1 normalized to β-actin are presented. miR-144-3p significantly repressed RXFP1 expression in donor lung fibroblasts in a dose dependent fashion (data analyzed by one-way ANOVA, p-value for trend 0.0006, R^2^=0.640). (*H*) Densitometry of α-SMA were normalized to β-actin is presented. α-SMA levels increased in a dose-dependent fashion (by one-way ANOVA, p-value for trend 0.0168, R^2^=0.51).

We further determined if the effects of miR-144-3p on the RXFP1 were detectable at protein level. Using western blot and densitometry analyses, we found that forced expression of miR-144-3p mimic in donor and IPF lung fibroblasts for 72 hours resulted in a significant repression of RXFP1 protein levels (Fig 2*C*). Extremely significant effects of tissue origin (IPF versus donor, *p*=0.0006, by two-way ANOVA, N=3) and miR-144-3p transfection (*p*=0.0004, by two-way ANOVA, N=3) on RXFP1 expression were detected. A significant interaction between tissue origin and miR-144-3p expression was also detected (p<0.03) (Fig 2*D*).

We previously found that the loss of RXFP1 is also associated with increased expression of the myofibroblast differentiation marker, α-SMA. To determine if miR-144-3p-mediated loss of RXFP1 is associated with α-SMA expression, we further analyzed its role in the response to miR-144-3p overexpression. We transfected donor and IPF lung fibroblasts with miR-144-3p or negative control mimic. Western blot and densitometry analysis showed that miR-144-3p overexpression resulted in increased levels of α-SMA in donor lung fibroblasts; however, α-SMA levels increased only slightly in IPF lung fibroblasts over baseline. Significant effects of tissue origin (IPF *versus* Donor, p=0.0038, by twoway ANOVA, N=3) on α-SMA expression were detected (Fig 2*E*). No significant interaction was detected.

Next, we determined if RXFP1 protein levels can be inhibited in a dose-dependent manner in donor fibroblasts using a range of mimic concentrations (0, 0.1nM, 1nM, 5nM and 10 nM). Western blot and densitometry analysis show that miR-144-3p mimic decreased RXFP1 with increasing concentrations of miR-144-3p mimic while completely inhibiting expression of RXFP1 at a concentration of 10 nM in donor lung fibroblasts (Fig 2*F*). Significant effects of concentration of miR-144-3p treatment on RXFP1 expression were detected (by one-way ANOVA, p-value for trend 0.0006, R^2^=0.64) (Fig 2*G*). While RXFP1 levels decreased, α-SMA levels increased in a dose-dependent manner (by one-way ANOVA, p-value for trend 0.0168, R^2^=0.51) (Fig 2*H*).

*Anti-miR-144-3p reverses the suppression of RXFP1 in IPF lung fibroblasts*—Having shown that RXFP1 is a targeted gene of miR-144-3p, we next examined whether forced expression of the miR-144-3p inhibitor (antagomiR) in donor and IPF lung fibroblasts would increase RXFP1 expression. As shown in Fig 3*A*, western blot and densitometry analyses show that the expression of RXFP1 protein was significantly increased by forced expression of the miR-144-3p antagomiR in IPF lung fibroblasts compared to donor lung fibroblasts. Significant effects of tissue origin (IPF *versus* Donor, p=0.007, by two-way ANOVA, N=3) and miR-144-3p antagomiR treatment (p<0.007, by two-way ANOVA, N=3) on RXFP1 expression were detected (Fig 3*B*). No significant interaction was detected.

**FIGURE 3.**
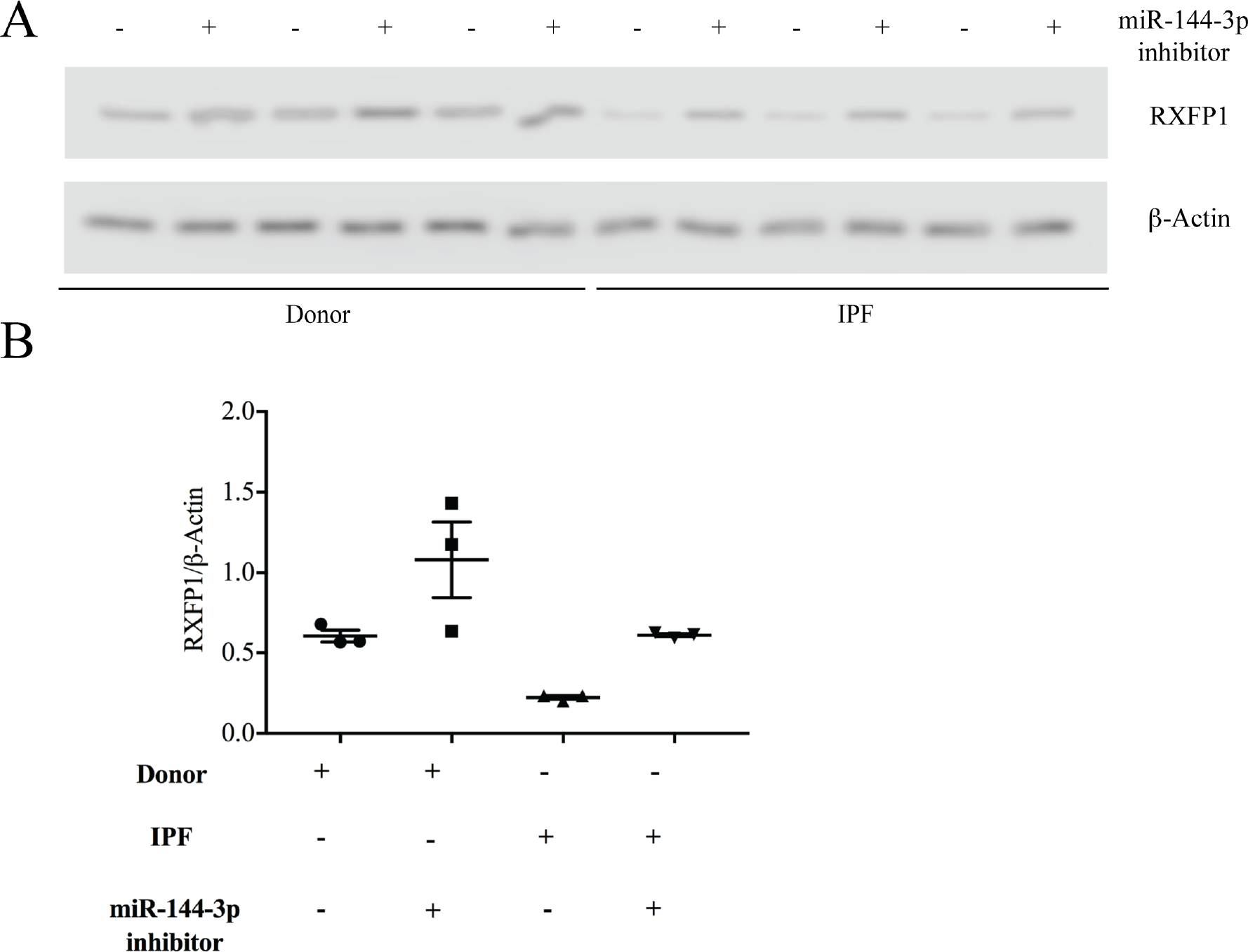
miR-144-3p antagomiR reversed the suppression of RXFP1 in IPF lung fibroblasts. (A) Donor and IPF lung fibroblasts were transfected with 50 nM miR-144-3p antagomiR or control. miR-144-3p antagomiR significantly increased expression of RXFP1 in IPF lung fibroblasts compared to donor lung fibroblasts. (*B*) Image Studio^™^ quantification of band intensity of RXFP1 was normalized to β-actin. Significant effects of tissue origin (IPF *v* Donor, p=0.007, by two-way ANOVA, N=3) and miR-144-3p antagomiR treatment (p<0.007, by two-way ANOVA, N=3) on RXFP1 expression were detected. No significant interaction was detected.

*miR-144-3p directly targets the 3’UTR region of RXFP1 mRNA—*We cloned the 3’-UTR portion of the RXFP1 containing the putative miR-144-3p binding sequence that is complementary to the miR-144-3p seed sequence in pmirGLO luciferase reporter plasmid (Fig 4*A*). In a parallel experiment, the 3’ UTR of RXFP1 complementary to the miR-144-3p seed sequence was mutated and cloned in the same reporter plasmid. Transient transfection of HEK293T cells with the RXFP1-3’-UTR construct along with miR-144-3p mimic (100 nM) led to a significant decrease (∼50%) in reporter expression when compared with the empty control vector or with scrambled control mimic (Fig 4*B*). The luciferase activity of the pmirGLO vector containing the 3’ UTR of RXFP1 with a mutated miR-144-3p target site was unaffected by simultaneous transfection with miR-144-3p mimic. These results suggest that miR-144-3p binds to the 3’UTR of RXFP1 to affect mRNA stability. To further verify direct targeting in primary human lung fibroblasts, lentiviral constructs carrying the complete RXFP1 3’UTR or miR-144-3p target site mutated 3’UTR sequences were used to transduce donor and IPF fibroblasts. Lung fibroblasts infected with wild type RXFP1 3’UTR construct showed reduced luciferase activity. Those infected with the mutant RXFP1 3’UTR construct showed diminished inhibition of luciferase activity (Fig 4*C*). These data indicate that endogenous miR-144-3p directly targets RXFP1 and regulates its expression in IPF patient lung fibroblasts.

**FIGURE 4.**
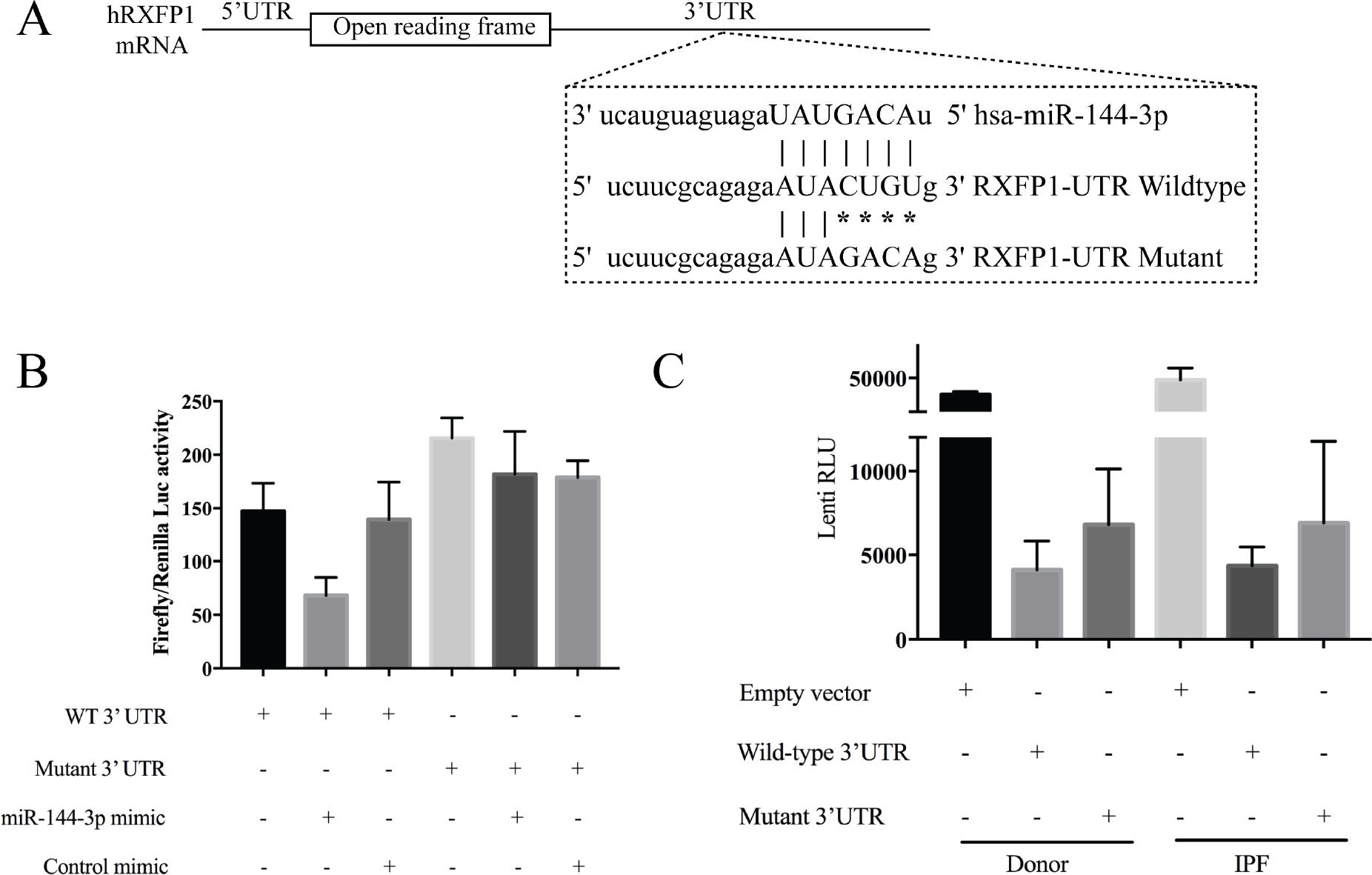
miR-144-3p directly targets the 3’UTR region of RXFP1 mRNA. (*A*) Depiction of pmirGLO dual-luciferase reporter construct for WT 3’UTR region of RXFP1 seed region and the mutation (*B*) 293T cells were plated 24 hours prior to transfection. Cells were then cotransfected with 50 ng of either pmirGLO vector carrying WT 3’ UTR of RXFP1 or mutated 3’ UTR of RXFP1 with and without 100 nM miR-144-3p mimic or control mimic. miR-144-3p significantly repressed the luciferase activity of the reporter containing the WT 3’ UTR of RXFP1, whereas mutated construct was insensitive in 293T cells. Luciferase activity was measured using *Renilla* luciferase as an internal control. Mean ± S.D, N=3 (*C*) When a lentiviral luciferase reporter construct carrying WT 3’ UTR of RXFP1 was used to transduce both Donor and IPF lung fibroblasts, there was a reduction in luciferase activity compared to mutated 3’UTR of RXFP1. Protein concentrations were measured using Pierce^™^ BCA protein kit assay and were used to normalize luciferase values. Mean ± S.D, N=3.

## DISCUSSION

In this study, we have shown increased expression of miR-144-3p in primary lung fibroblasts derived from IPF patients. Increased expression of miR-144-3p in donor lung fibroblasts, that contain significantly less miR-144-3p at baseline, led to decreased expression of RXFP1. We also show that blockade of miR-144-3p with an antagonist increased RXFP1 expression in IPF fibroblasts. Finally, we have shown that RXFP1 3’UTR luciferase reporter activity is decreased in IPF fibroblasts and that luciferase activity is decreased to a lesser extent containing a mutated miR-144-3p binding site. Together, these data suggest that increased miR-144-3p expression in IPF fibroblasts may, at least in part, explain decreased expression of RXFP1 in IPF fibroblasts.

A previous report has shown that the levels of miR-144-3p are elevated in the biopsy specimens from patients with the bronchiolitis obliterans syndrome (BOS)—a form of airway fibrosis associated with chronic rejection following lung transplant. In this study, forced expression of miR-144-3p mimic increased levels of TGFβ and *vice versa*, which increased α-SMA and F-actin levels in MRC5 fibroblasts (21). This, perhaps, indicates a feed-forward loop between miR-144-3p and TGFβ, that amplifies TGFβ signaling. However, in the present study TGFβ had little impact on expression of miR-144-3p, at least in IPF lung fibroblasts. Elevated expression of miR-144-3p in IPF lung fibroblasts may partially explain this observation. Therefore, we suggest that increased levels of miR-144-3p in IPF fibroblasts may be regulated by a non-TGFβ mediated pathway. Several previous reports have indicated that miR-144-3p expression is regulated by various stimuli and pathways. Hypoxia has been shown to decrease levels of miR-144-3p in PC3 prostate cancer cells (22). The mechanism underlying increased expression of miR-144-3p levels in IPF fibroblasts is unclear and needs further investigation.

MiR-144-3p is the mature microRNA excised from the precursor miR-144 that is encoded by the miR-144/451 gene cluster on chromosome 17q. Little is known about miR-144-3p in fibrosis, but the precursor microRNA is associated with reduced expression of the Smad signaling repressor transforming growth factor beta (TGFβ)-induced factor homeobox (TGIF1) (21). The transcription factors GATA1 (23) and GATA4 are thought to positively regulate miR-144 expression in mice (24). PAX4 has also been shown to negatively regulate the miR-144/451 cluster in human epithelial cancer metastasis (25). Another recent report has shown that promoter region of miR-144/451 cluster is composed of 6 potential AP1/C-Jun binding sites with a positive regulatory role in controlling the cluster in human neuroblastoma SH-SY5Y cells (26). Expression of miR-144 has been previously reported to increase during erythropoiesis in mice and humans (27). Expression of the cluster of miR-144/451 has been shown to be significantly low in non-small cell lung cancers (NSCLCs) compared to adjacent normal lung tissue and is also associated with inhibition of cancer cell migration and invasion (28).

In our present study we show that levels of α-SMA increased dose-dependently with increasing concentrations of miR-144-3p mimic in lung fibroblasts (Fig 2*B*). These results are consistent with our previous report where we showed that myofibroblast function is critically linked to RXFP1 expression (7). We and others have previously shown that Relaxin (7,29,30) or Relaxin-like agonists blocked the expression of α-SMA, an effect which is lost in the absence of RXFP1. Relaxin has been shown to inhibit myofibroblast function by inactivating Rho-ROCK pathway (31). Increased expression of α-SMA in miR-144-3p-transfected cells may reflect the lack of relaxin signaling following the loss of RXFP1.

The regulation of RXFP1 in other tissues and other cells may be affected by other microRNAs. The WT lentiviral construct carried the entire 3’ UTR sequence of RXFP1, which is predicted to have other potential microRNA target sites in addition to the miR-144-3p site. The mutant construct carried the miR-144-3p mutation only leaving the other target sites intact. We suggest that other unaltered microRNA target sites may explain the mildly-diminished inhibition of luciferase activity in the miR-144-3p mutant transduction experiment compared to the WT construct. Future study is needed to understand the regulation of RXFP1 expression in other tissues and other cells.

In conclusion, we have identified a potential mechanism for regulating expression of RXFP1 in pulmonary fibrosis. Because of the importance of relaxin signaling to fibrosis, we speculate that molecular strategies to increase RXFP1 expression may potentially sensitize fibroblasts, the principal effector cells of fibrosis, to reap the beneficial effects of relaxin. Further study is needed to consider miR-144-3p inhibitors in animal models of fibrosis.

## EXPERIMENTAL PROCEDURES

This study was conducted in accordance with University of Pittsburgh IRB protocol # PRO14120072.

*Bioinformatics*—Prediction of miRNA targets was conducted using miRanda (http://www.microrna.org/) (32) and Targetscan 7.0 database (http://www.targetscan.org/vert71/) (33). The putative sequences of primary microRNAs and the 3’UTR of RXFP1, 2 and 3 were retrieved from the National Center for Biotechnology Information (NCBI) (http://www.ncbi.nlm.nih.gov) and ENSEMBLE server (http://www.ensemble.org) (34) and miRBase (http://www.mirbase.org) (35).

*Primary cell culture*—Primary lung fibroblasts were obtained from Dr. Mauricio Rojas (36). Donor human fibroblasts were isolated from lungs that appeared to have no injury by histology but were deemed unacceptable for lung transplant. IPF lung fibroblasts were obtained from patients either at explant or at autopsy (37). All fibroblasts were maintained in Dulbecco’s modified Eagle’s medium supplemented with 10% fetal bovine serum and were used between passages 3 and 6 and used for expression analysis of both RXFP1 and miR-144-3p. MiR-144 antagomiR (Dharmacon) or MirVana miR-144 mimic (Ambion) were used to modulate miR144 levels. RXFP1 protein expression was analyzed using Western blot.

*RNA extraction*—Briefly, total RNAs were extracted from the donor and IPF lung fibroblasts using RNeasy (Qiagen). The RNA samples were reverse transcribed using the High Capacity cDNA Reverse Transcription Kit (Applied Biosystems, Foster City, CA). The product from each reverse transcription reaction was pre-amplified and then the mRNA expression analysis was done by qRT-PCR using ABI TaqMan system (Life Technologies) following the manufacturer’s recommended protocol. qRT-PCR primers were purchased from Qiagen (Hsa-PPIA Cat # QT01669542, Hsa-RXFP1 Cat # QT00041720, Hsa-RXFP2 Cat # 00095725, Hsa-RXFP3 Cat # 00210133). PPIA [peptidylprolyl isomerase A (cyclophilin A)] was used as the housekeeping gene for normalization. The global normalization process included the subtraction of the mean C_T_ value of the reference set from the C_T_ value of each gene of the same sample. Quantification of each sample is shown as 2^-ΔΔCt^ values.

*TaqMan Real-time RT-PCR for miRNAs*—Small RNAs were isolated from cells following the protocol of the miRNeasy kit (Qiagen). 100-300 ng of RNA sample was reverse-transcribed into cDNA using the TaqMan™ Advanced miRNA cDNA Synthesis Kit (Applied Biosystems). Taqman probes (TaqMan^™^ microRNA Control Assay for RNU43 and TaqMan^™^ microRNA Assay for hsa-miR-144) were purchased from Applied Biosystems. Real-time quantitative PCR was performed using the TaqMan Fast Advanced Master Mix (Applied Biosystems). Quantification of each sample is shown as 2^-ΔΔCt^ values.

*Western Blotting*—Western blotting was performed as described previously (7). Anti-RXFP1 (ab72159) and Anti-aSMA antibodies (ab7817) were purchased from Abcam. Mouse anti-actin (sc-47778) was from Santa Cruz Biotechnology.

*Design of reporter constructs and Luciferase assays*—Duplexed oligonucleotide pairs (IDT) were designed to contain the predicted miR-144-3p binding region in the RXFP1 3’UTR and when annealed and ligated into the pmirGLO vector, resulted in the miR-144-3p target region in the correct 5’ to 3’ orientation. The sequences of the duplexes used were as follows: RXFP1-Mir144-WT Duplex (Sense: 5’ AAACTA**GCGGCCGC**TAGTTCTTCGCAGA GA<u>*ATACTGT*</u>GGGGGTGT; Antisense: 5’ *CTAGACACCCCC<u>*ACAGTAT*</u>TCTCTGCGA* AGAACTA**GCGGCCGC** TAGTTT); RXFP1-Mir144-Mismatch Duplex (Sense: 5’ AAACTA**GCGGCCGC** TAGTTCTTCGCAGA GA<u>*ATAAAT*</u>GGGGGTGT, Antisense: 5’ CT AGACACCCCC<u>*ATTTAT*</u>TCTCTGCGAA GAACTA**GCGGCCGC** TAGTTT). Overhangs in the designed duplex were complementary to those generated by *PmeI* and *XbaI* double digestion of the pmirGLO Vector (Promega). *Not1* internal restriction site was designed into the oligo duplex for clone confirmation. pmirGLO Vector was linearized with *Pme1* and *Xba1* to generate overhangs that are complementary to the duplex overhangs. 4ng of duplexed oligonucleotide and 50 ng of linearized vector were ligated using a standard ligation protocol. Ligated pmirGLO was transformed using high-efficiency TOP10 competent cells (NEB). Clones were selected on carbenicillin-containing plates, and then screened for clones containing the duplex by digesting miniprep-purified DNA using the internal *NotI* site. All constructs and mutants were verified by DNA sequencing at the Genomics core facility at University of Pittsburgh. The purified plasmid DNA was used directly in transfections. The resulting constructs were designated as pmirGLO-RXFP1-WT and pmirGLO-RXFP1-MsM. HEK-293T cells in a 24-well plate were transfected with 50 ng of pmirGLo-RXFPI-WT in triplicates and 100 nM control mimic or 100 nM miR-144-3p mimic (Ambion) using Lipofectamine 2000 reagent (Invitrogen). 24 hrs post-transfection, luciferase assay was performed on the lysates from the cells using a Dual-Luciferase Assay System (E1910, Promega) on Spectramax L instrument. Data were normalized by ratio of firefly and renilla luciferase activities.

*Lentiviral vector transduction in lung fibroblasts*—Lentiviral vectors carrying the 3’ UTR region of RXFP1, pLenti-RXFP1-UTR-Luc and its mutant, pLenti-UTR-GFP-Blank, and pLenti-UTR-Luc-Blank were purchased from applied biological materials Inc. (abmgood. BC, Canada). Lentiviral packaging was performed using viral packaging plasmids from the ‘ViraPower’ kit (Invitrogen) according to the manufacturer’s instructions. Filtered viral supernatants were obtained from transfection of 293FT cells according to the manufacturer’s protocol. qPCR Lentivirus Titration Kit (abmgood) was used to measure the lentiviral titres according to the manufacturer’s instructions. Viral medium was added to complete media in presence of Polybrene (0.8 μg/ml) at a multiplicity of infection (MOI) of 5 to infect Donor and IPF lung fibroblasts for 72 hrs. Cells were lysed in passive lysis buffer and dual-luciferase assays were performed as described above. Protein concentrations were measured using Pierce BCA micro assay and were used to normalize luciferase values (RLU/μg; Relative light units/μg of protein).

*Transfection of miRNAs*— Fibroblasts were transfected with miRNA mimics or miRNA inhibitors using HiperFect transfection reagent (Qiagen). Control or miR-144-3p mimics were purchased from Invitrogen. Control or miR-144-3p inhibitors were purchased from Qiagen. Briefly, transfection complexes were generated by mixing 0-10 nM of miRNA mimic/ control mimic or 50 nM inhibitor or control inhibitor, 100 μL of DMEM media without serum, and 6μL of HiPerFect Transfection Reagent. After 10-minute incubation at room temperature, the transfection complexes were added drop-wise onto 2 X 10^5^ cells per well of a 6-well plate in 2000 μL of culture medium with 1% FBS but without antibiotics. Cells were transfected on the day of plating and again on the next day.

*Statistical Analysis*—Analysis of variance (ANOVA) was performed for multiple group comparisons. p<0.05 was considered statistically significant. Statistical testing is indicated in the text and the figure legends. Data were analyzed using GraphPad Prism7 software.

## Acknowledgements

None

## Author Contributions

JT, HB, JAD, and SBS conducted experiments and analyzed the results. JT, JAD, SBS conducted qRT-PCR and western blot experiments. HB conducted the luciferase experiments. DJK, JT, and YZ conceived and planned experiments. HB, JT, JAD, SBS, YZ and DJK wrote the manuscript.

The abbreviations used are: RXFP1, relaxin/insulin-like family peptide receptor 1; TGF, transforming growth factor; DMEM, Dulbecco’s modified Eagle’s medium; GPCR, G-protein coupled receptor; ANOVA, analysis of variance; IPF, Idiopathic pulmonary fibrosis; SSc, Scleroderma.

